# Selecting methods for draft GEM generation in multicellular eukaryotes: a comparative analysis

**DOI:** 10.1101/2025.06.23.661012

**Authors:** Natalia E. Jiménez, Mikael Espinoza, Sebastián Mejías, Sebastián Mendoza, Ignacia Segovia, J. C. Salgado, Carlos Conca, Ziomara. P. Gerdtzen

## Abstract

Motivated by multiple strategies that have successfully implemented genome-scale models (GEMs) into their pipeline, several approaches have been developed for automatic generation of draft GEMs. However, most of these methods are not optimized for their use for multicellular eukaryotes and their performance for this task is unclear. In this work we present a comparative analysis of seven automated reconstruction tools (AuReMe, carveMe, Merlin, modelSEED, Pathway tools, Raven and Reconstructor) applied to three multicellular eukaryotes: the mosquito *Aedes aegypti*, the CHO (Chinese Hamster Ovary) cell line from *Cricetulus griseus* and the brown algae *Ectocarpus siliculosus*. Evaluation of these tools was based on metrics for network size, functionality, consistency, representation of organelle-specific functions and organism-specific metabolites, annotation quality and execution time. Finding that similarity of obtained metabolic networks is highly influenced by databases in which these methods base their predictions over phylogeny. Our works aims at providing a practical resource to guide researchers in selecting methods for draft generation tailored to organism characteristics and research goals.

**Author summary:** Genome-scale models (GEMs) represent all the potential biochemical transformations that a specific organism can carry out based on the information encoded in its genome. Although these models are a powerful tool for analyzing omics datasets and simulating the metabolic effects of genetic modifications or changing media composition, the process of manually reconstructing a genome-scale model is complex and time-consuming. Motivated by their potential applications, several tools have been developed for the automated generation of draft GEMs, however, most of them are oriented to simpler organisms such as bacteria or single-cell eukaryotes, while their relative performance for modeling multicellular eukaryotes is unclear. In this work, we compared seven tools for draft GEMs reconstruction of three organisms: for the mosquito *Aedes aegypti*, the brown algae *Ectocarpus siliculosus* and the CHO cell line from *Cricetulus griseus.* Our results showed that no tool systematically outperformed others, suggesting that method selection is influenced by several factors such as organism-specific data availability and their intended application.

## Background

With the reduction of DNA sequencing costs (1–4), genome-scale models (GEMs) have emerged as a powerful tool for biological discovery and metabolic engineering, due to their ability to simulate gene knockouts (5–8), describe multi-species relationships (9,10), contextualize omics data (11,12), identify gene targets for cancer therapy (13,14), and provide a framework for mechanistic understanding of phenotypes of interest (15). These models provide a global representation of all biochemical transformations that could be carried out by a specific organism and the relationship between these transformations and their genes, given by logical rules called Gene Protein Reaction (GPR) associations (16,17).

Due to the multiple frameworks that have successfully integrated GEMs into their pipeline (18–20), motivation for development of new metabolic reconstructions has increased, and the inclusion of more complex organisms is impending. However, the process of manually generating GEMs is complex and time-consuming, often taking up to several years (17). Hence, several automatic strategies have been developed to generate draft GEMs from sequenced genomes (21), which rely on specialized biochemical databases (22–24), their own genome annotation (25) or a curated available version including EC numbers to establish a draft version of the desired GEM (26–30). Orthology inferences can also be used to retrieve candidate reactions (31) and information from previously curated models can aid in this process (22,24,31).

While these automated approaches have been successfully applied for generating draft GEMs, their primary focus has been on prokaryotes and unicellular eukaryotes such as *S. cerevisiae* (32) and *Y. lipolytica* (33). There are fewer examples of their use in multicellular eukaryotes, with some exceptions such as plantSEED (based on modelSEED) and the use of these strategies for generating plant models (25,29), macroalgae (34) and mammalian cell lines (35). Relative performances of different automated reconstruction tools have not been systematically assessed for multicellular eukaryotes. Furthermore it is not clear which model metrics would be relevant to compare in order to benefit draft GEM applications. This is critical as metabolic understanding of these organisms have deep implications for public health (36), sustainable economy (34) and the biopharmaceutical industry (35).

In this work, we provide a perspective on the automatic draft GEM reconstruction methods for multicellular eukaryotic organisms. Three organisms with different degrees of prior metabolic characterization were chosen for this comparison: (i) the Chinese Hamster Ovary (CHO) cell line from *Cricetulus griseus*, which has an extensively curated GEM widely used for basic and applied metabolic research (37–39), (ii) the brown algae *Ectocarpus siliculosus* whose interactions with its microbial communities have been studied using GEMs (40), and (iii) the mosquito *Aedes aegypti,* which has a reconstruction used for studying its interactions with its endosymbiont *Wolbachia pipientis* (36,41). In this study, we provide a thorough analysis of the most relevant automated tools for generation of draft GEMs compatible with eukaryotic organisms, highlighting relevant challenges that arise when working with such organisms.

## Results

### Metric selection

Eukaryotic organisms are complex, they harbor specialized metabolic pathways yet to be discovered, perform chemical transformations in organelles, and exhibit higher genome size with respect to prokaryotes, posing additional challenges for generating GEMs. An ideal method should be able to overcome these challenges and provide a draft GEM that includes prior biological knowledge from the organism of interest, represents multiple compartments whilst preserving association with organelle-specific enzymes, and that is able to synthesize as many metabolites that comprise its biomass as possible. Generated models should also include links to different databases for metabolites and enzymes to facilitate the integration with models from other databases as well as from omic data analyses (42).

In this work we analyzed seven methods for generation of draft GEMs: AuReMe (31), carveMe (27), merlin (25), plantSEED (29), Pathway Tools (26), Raven (24) and Reconstructor (22). Ten metrics were selected to characterize these methods, aligned with the desired features described previously. Selected metrics reflect the size of draft GEMs (number of reactions, metabolites, genes and compartments), how close they are to be able to produce biomass (*i.e.* be *functional*) (biomass metabolites, flux reactions, consistency), how fast is the reconstruction process (time), as well as its quality for their use in applications such as analyzing gene expression data (annotation score, enzymatic reactions) (**Figure 1, Table S1**).

**Figure 1:**
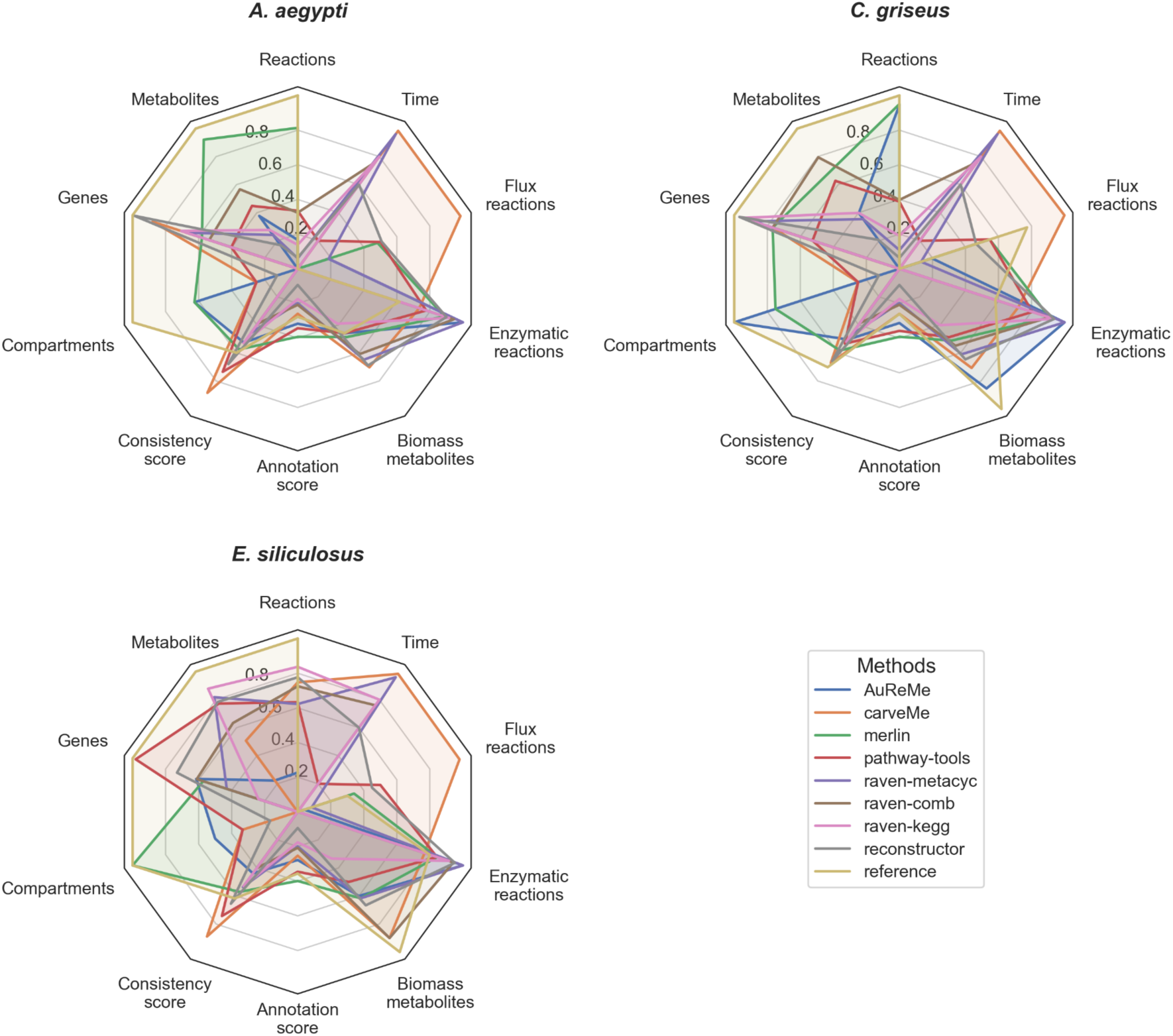
Obtained draft models for *A. aegypti*, *C. griseus* and *E. siliculosus*, exhibit differences regarding their size (reactions, metabolites, genes, compartments), how close they are to be functional (flux reactions, biomass metabolites), and how well annotated they are (annotation score, enzymatic reactions). Metrics are normalized in regards to their reference model.

### Obtained drafts are influenced by design decisions of automated reconstruction tools

Figure 1 shows results obtained for *A. aegypti*, *C. griseus*, and *E. siliculosus*, depicting that results are influenced by the target organism as well as decisions made in the design of such methods. In particular, for plantSEED (**Table S1**), all generated models were identical in terms of metabolites and reactions (1,179 and 1,121, respectively) and differ only in the number of genes (1 (Unknown) *A. aegypti*, 634 *C. griseus*, 728 *E. siliculosus*), reflecting an intensive gap filling (64% of reactions are able to carry flux) performed by default by this platform (29) (**Table S2**).

Reconstructor (22), also based in the modelSEED database, yields better results to those obtained by plantSEED, whilst preserving the advantages that this highly curated database presents. Although it is based in bacteria, obtained models yield on average a larger number of genes (128%), reactions (20%) and metabolites (77%) compared to other bacterial-based methods like carveMe. However, its annotation score is considerably low (less than 0.1) being the worst in this category, which would be a disadvantage for analysis of omic datasets, requiring extra mapping efforts.

carveMe generates models with the highest proportion of reactions being able to carry flux (98-99%), a consequence of its top-down approach for draft generation. This method starts its reconstruction process from an universal model and it eliminates reactions for which there is no evidence of its presence (27).

AuReMe (31) incorporates information from a template model to retrieve additional reactions and associated genes based on orthology. Criteria for model selection is based on phylogenetic closeness, with the exception of *E. siliculosus*, for which a model for *S. japonica* (43) was excluded since it was based on the curated model of *E. siliculosus* (34) and a *Chlamydomonas reinhardtii* model was used instead (44); for *A. aegypti* a *Drosophila melanogaster* model was used as a template, and for *C. griseus* a *Mus musculus* model (45) was selected to this end. Obtained draft models are highly influenced by characteristics of their corresponding templates, where inclusion of reactions and metabolites is driven by how curated each model is, rather than phylogeny alone, where bigger models that include more metabolic information (such as *H. sapiens* (46) and *S. japonica* (47)) resulting on bigger models **(Table S1)**.

Models generated using Merlin include predictions of subcellular localization of enzymes codified in their genome (deeploc2 (48)). As a consequence, models generated by this tool exhibit more compartments in average than others (merlin: 7, carveMe: 3, reconstructor: 2, raven: 1, Pathway tools: 3, AuReMe: 6-10); thus leading to having more reactions (2,894 *A. aegypti*, 6,390 *C. griseus*, 3,813 *E. siliculosus*). Particularly for *C. griseus*, 57% of their 5,808 metabolites are duplicated in different cellular compartments. These models also exhibit high variability in reactions that can carry flux (13 to 54%) showing an organism-dependent behavior in regards to their functionality.

Pathway tools generated models also exhibit an organism-dependent behavior, displaying higher gene numbers in *A. aegypti* (5,509) compared to *C. griseus* (3,891), despite being metabolic networks with comparable size in terms of metabolites and reactions. *E. siliculosus* exhibits more reactions than genes, which could be due to its automated gap filling, and also to biological properties to this macroalgae which is unable to grow axenically (26). This tool also includes generic reactions that have to be furtherly curated by the user of the model, and that can generate issues in Flux Balance Analysis simulations for polymerization reactions if ignored.

Raven 2 (Wang et al., 2018) allows to generate draft GEMs using different approaches, a Metacyc-based draft (‘raven-metacyc’ in this work) generated from amino acid fasta files. A KEGG-based draft based on sequence homology (‘raven-kegg’), as well as a combination of both approaches (‘raven-comb’). Raven represents a single compartment and uses a conservative approach where only reactions with gene associations are included. As a consequence of the resulting metabolic gaps, these models are characterized by a lower proportion of reactions able to carry flux (8% *E. siliculosus* to 18% *A. aegypti*). Particularly, in models generated using Raven linked with the KEGG database, no reactions are able to carry flux in all models due to the lack of exchange reactions.

Time required to obtain draft reconstructions varied greatly across evaluated tools, ranging from minutes to days. Merlin had the longest processing time with draft reconstruction taking up to five days (**Table S1**). This delay is primarily due to the BLAST performance on their servers as well as the use of external applications such as deeploc2 and the constant input from the user is required and also contributes to longer processing time. On the other hand, carveMe was found to have the shortest processing time taking up to 5 minutes, whilst AuReMe exhibited changes in time depending on the selected template, ranging from 20 minutes to 5 hours.

Size of the obtained metabolic networks (number of reactions and metabolites) is highly influenced by information available on databases used by each method. This is more evident in the case of *C. griseus* models, which being closer to *H. sapiens* are reconstructed using information from this organism yielding models with over 6,000 reactions (6,968 AuReMe, 6,390 merlin), where *A. aegypti* and *E. siliculosus* exhibit values near 3,000 reactions (3,813 merlin, 3,403 AuReMe).

### Organism-dependent functionality analysis hints issues with boundary reaction definition in draft models

Given the relevance of biomass synthesis in GEMs (49), draft models were studied in terms of how closely they represent organism-specific curated versions of their biomass, rather than proposed biomass reactions from their respective methods. To achieve this, presence and producibility of biomass metabolites from reference models (**Figure S3**) was assessed in obtained drafts (Figure 2), finding both organism and method-dependent issues in the functionality of these models.

**Figure 2:**
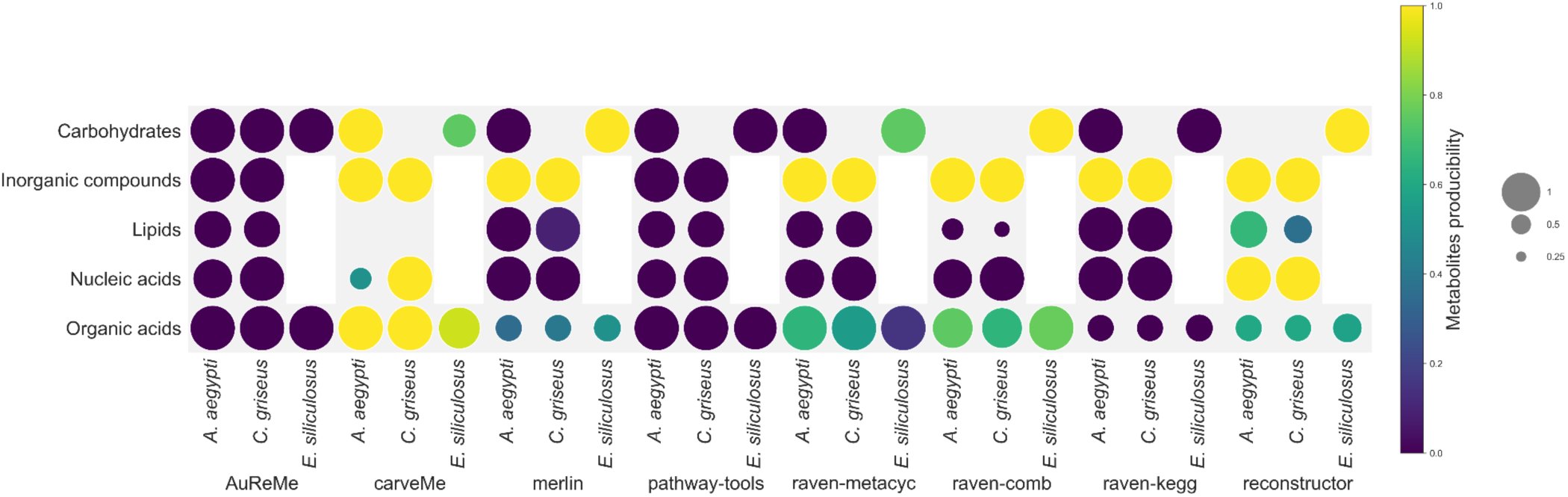
Presence and producibility of organism-specific biomass metabolites for obtained draft models. Fraction of present metabolites of a given category is depicted by circle size. Fraction of present metabolites that are also producible using complete media is depicted by circle color. White rectangles indicate categories that are not present in their respective reference organism-specific biomass.

*E. siliculosus* curated GEM presents a simpler representation of its biomass comprising organic acids (mostly amino acids, together with carboxylic acids) and carbohydrates (glycerol, mannitol, glycollate, glycerate and threo-ds-iso-citrate) (34), whilst *A. aegypti* and *C. griseus* exhibit more complex biomass representations including lipids, nucleic acids and inorganic compounds (**Figure S3**).

Inorganic compounds are ions and metals that are directly imported into the cell from their surrounding media, their lack of producibility on certain methods (AuReMe, Pathway tools) hints issues with the definition of exchange reactions in draft models. These reactions represent boundary conditions of the surroundings of the modeled organisms. In fact, despite including metabolites from their biomass for all three organisms, models generated with Pathway tools are unable to produce them in complete media, for which manual curation is required to obtain functional models with this approach.

On the other hand, draft models obtained using Raven-metacyc, Raven-kegg, Raven-comb and Reconstructor are able to import inorganic compounds, and hence integrate them to their biomass. Reconstructor models are also able to produce nearly 60% of the organic acids present on their drafts, and together with carveMe are the only ones able to synthesize nucleic acids in complete media. Raven-metacyc and Raven-comb exhibit producibility of organic acids, but not for the remaining analyzed categories, hence not retrieving functional models.

Consistent with the top-down approach performed by carveMe, models generated with this tool exhibit the highest producibility of biomass metabolites. These draft models exhibit a reduced capability of production for nucleic acids in *A. aegypti* (50%, 4) and carbohydrates for *E. siliculosus* (75%, 3). However, no lipids were mapped into these draft models of the 6 and 11 present in *A. aegypti* and *C. griseus* biomass respectively, showing a potential database dependency on the presence of metabolites included in obtained draft models using these tools.

### Gene-compartment specificity reflects database information for different organisms

Eukaryotic organisms are highly compartmentalized thus having specific functions that are encoded by different enzymes. This results in bigger metabolic networks which include multiple versions of reactions in different compartments that are usually linked to different genes in their GPRs. We aimed to assess how different frameworks for GEMs reconstruction are able to incorporate this information on their draft models (Figure 3). Reference models exhibit a mixed distribution between three alternatives: reactions that are duplicated without including a gene association rule, duplicated reactions with the same GPR and duplicated reactions with different rules in less proportion. Draft models obtained using Raven were excluded from this analysis since they lack compartmentalization (Figure 1).

**Figure 3:**
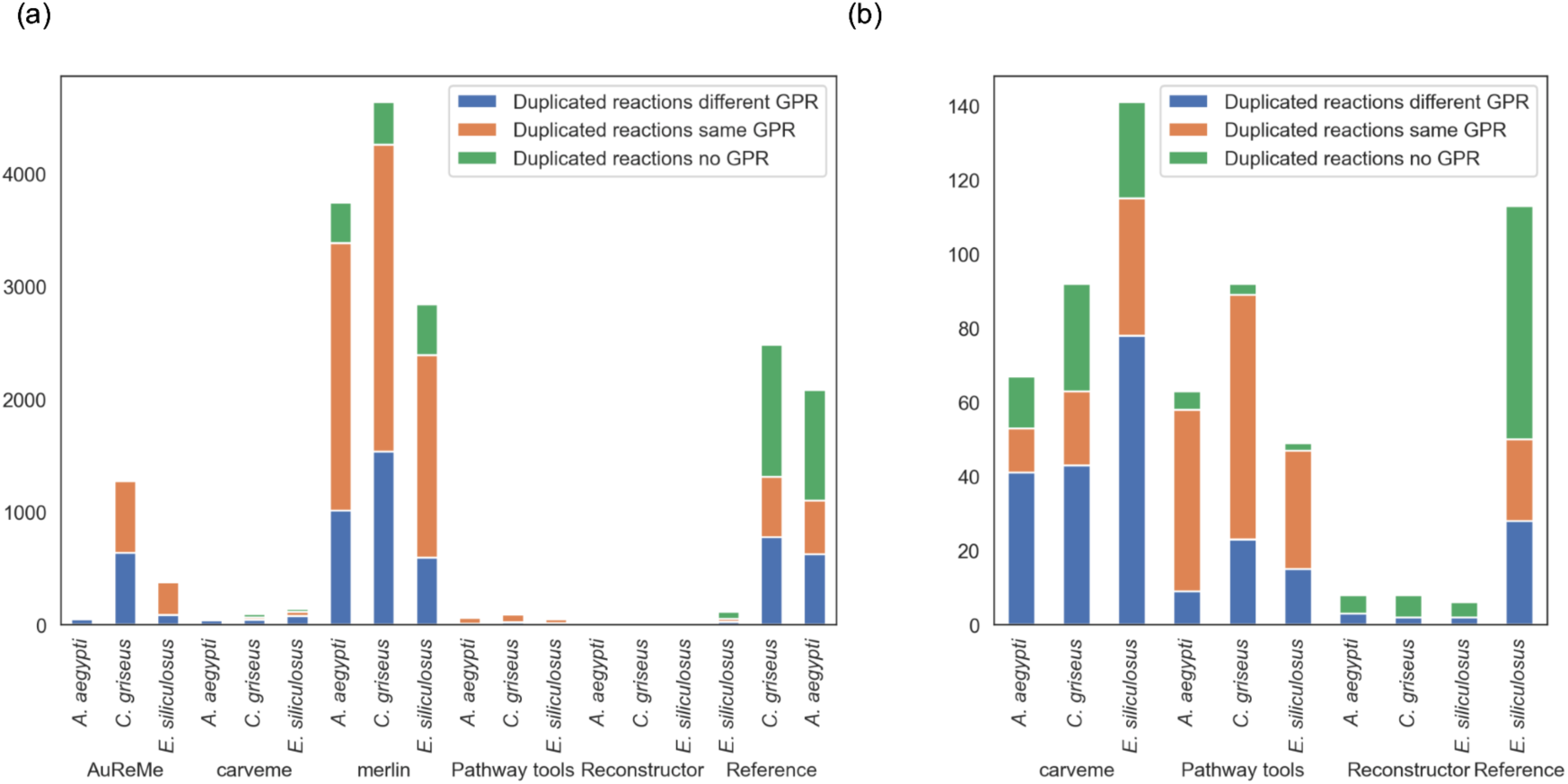
GPR assignment comparison between obtained draft genome-scale models. Duplicated reactions in different compartments were analyzed to check if they displayed gene associations, and if these are duplicated from reactions occurring in the cytoplasm. (a): all models, (b): models with smaller number of reactions for this analysis. au: AuReMe, cv: carveMe, me: merlin, pt: pathway tools, rec: reconstructor, ref: reference models.

Models generated using Merlin have been found to display higher numbers of reactions and compartments (*A. aegypti*: 2, *C. griseus* and *E. siliculosus*: 7) (Figure 1), however most of these are duplicates that share the same gene associations as their cytosolic counterparts (25% *A. aegypti*, 28% *C. griseus*, 30% *E. siliculosus*) (Figure 3a). Although AuReMe models display high numbers of compartments (*A. aegypti:* 6, *C. griseus*: 9, *E. siliculosus:* 10) the presence of duplicated reactions is lower than Merlin (*A. aegypti*: 3,742, *C. griseus*: 4,632, *E. siliculosus*: 2,840) and comparable to references models of their respective organisms.

Pathway Tools models exhibit low incidence of duplicated reactions, and most duplicates lack differences in their GPRs (Figure 3b). Similarly, carveMe includes three compartments (cytosol, extracellular, periplasm) and exhibits low presence of duplicates. Presence of bacterial compartments in draft models obtained with this tool requires additional curation of reactions associated with periplasm. Finally, models generated using Reconstructor exhibit the lowest number of duplicate reactions with no duplicated GPRs between compartments, as a consequence of the lower number of compartments exhibited in these models.

Reference models exhibit the greatest number of compartments (*A. aegypti* and *C. griseus*: 9, *E. siliculosus*: 7), and ratios of compartment specific genes of 19%, 39% and 12**%** for *A. aegypti*, *C. griseus* and *E. siliculosus* respectively, denoting an organism-specific availability of these information in databases that is also reflected in the obtained models, in particular for *C. griseus* for which information is improved by the highly characterized human metabolism.

### Representation of specialized metabolites is influenced by method rather than phylogeny

Multicellular organisms present specialized metabolites that are absent in prokaryotes, or unicellular eukaryotes. To depict the capability of draft models of representing such specialized metabolites, metabolomic sets were compiled for *A. aegypti* (50), CHO cells (35,51–59) and *E. siliculosus* (60), generating a list of query metabolites for these organisms (*A. aegypti*: 389, *C. griseus*: 306, *E. siliculosus*: 372) which were mapped to metaboAnalyst (61) retrieving a query list of metabolites for each organism whose presence was assessed on the obtained draft models for all organisms (*A. aegypti*: 54, *C. griseus*: 140, *E. siliculosus*: 217) (Figure 4a). 11 core metabolites were identified as being present in all datasets, which correspond to 7 amino acids and derivatives (methionine sulfoxide, proline, phenylalanine, histidine, ornithine, tryptophan, thymidine, arginine), 1 fatty acid (Pentadecanoic acid), 1 vitamin (Vitamin B5), and 1 nucleotide (AMP) (**S4 table**).

**Figure 4:**
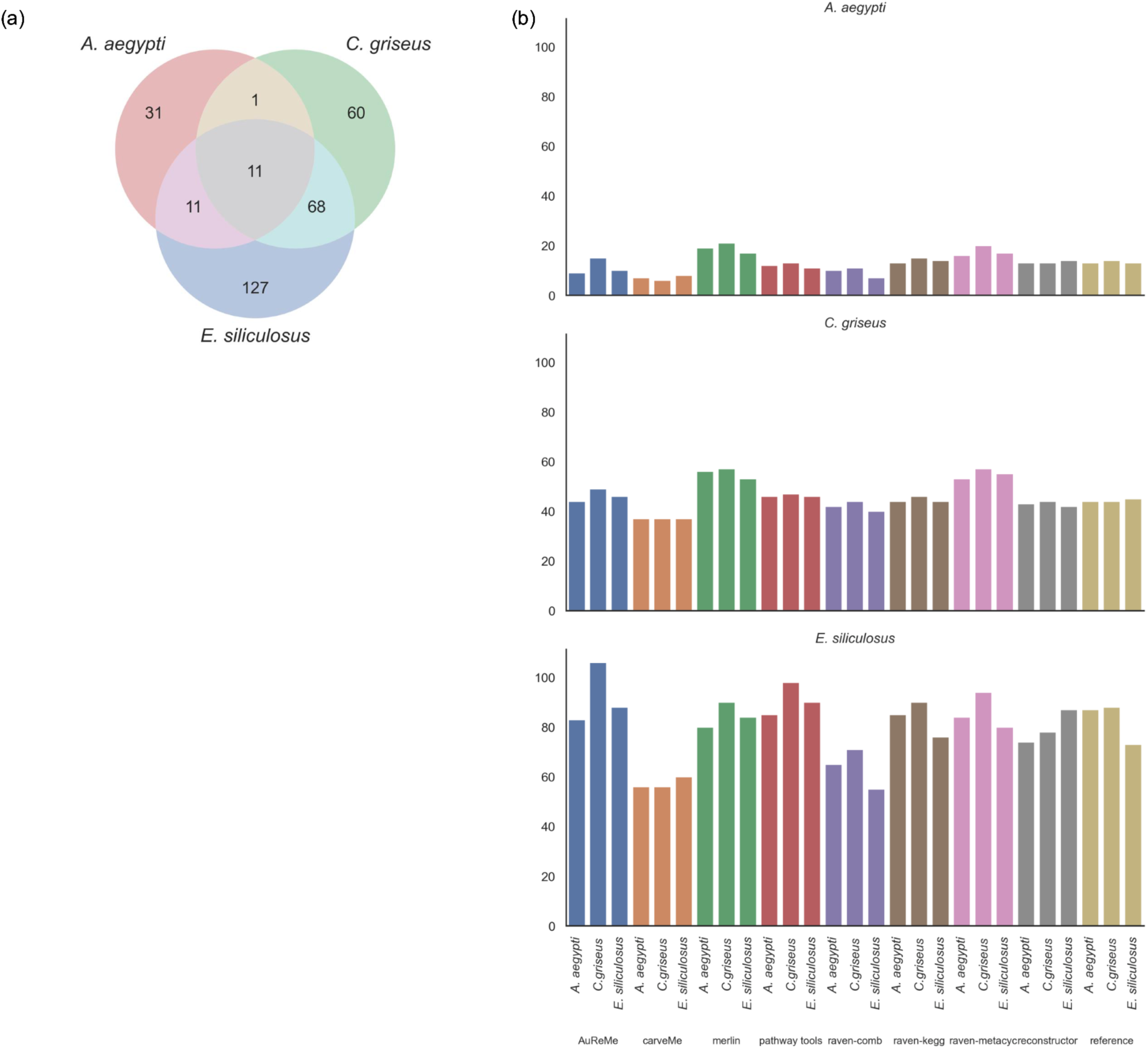
Presence of specialized metabolites in draft models. (a) Specialized metabolites were retrieved from literature for *A. aegypti, C. griseus* and *E. siliculosus*. (b) Lists of specialized metabolites were mapped to draft models of these organisms.

Results obtained from this mapping depend on: quality of the dataset that is being mapped, which organism is being modeled and which approach was used to generate such models. For the first point we see that for the *C. griseus* metabolome mapping carveMe and Reconstructor exhibit nearly identical metabolite mapping despite the modeled organism. This is consistent with the composition of this dataset, where only 43% metabolites are unique to this dataset in contrast with *A. aegypti* and *E. siliculosus* which include 57 and 58% unique metabolites respectively.

Obtained results hint a method dependency on specialized metabolite mapping. In fact, the number of mapped metabolites presents higher standard deviations when comparing drafts for the same organism generated using different tools (5.5-20) than for different organisms generated by the same approach (0–15) (Table S5). Mappings performed for models generated with AuReMe present the highest variation (standard deviation of 4.75 for *A. aegypti* metabolome mapping, 5.68 for *C. griseus* and 15.69 for *E. siliculosus*), highlighting the ability of this method to capture specificity displayed by models retrieved using these tools. Other methods are characterized by lower variations of mapping between different models, for example, carveMe models in *E. siliculosus* and *C. griseus* display almost identical results for all the modeled organisms. Furthermore, both method and organism were found to significantly influence metabolite coverage (two-way ANOVA test, p < 1e-9 and p < 1e-48, respectively) with a significant interaction effect between method and organism (p ≈ 6.3e-6) suggesting that specialized metabolite mapping performance varies depending both on the organism and method.

### Databases influence draft metabolic composition rather than phylogeny

Given our previous results we aim at determining which factor has a bigger influence on the structure of metabolic networks obtained using these tools. We studied the presence of metabolites and reactions, regardless of their associated compartment, finding that methods have a higher influence than phylogeny in how similar their metabolic networks are (Figure 5 **and Figure S7**). To achieve this, a binary vector was constructed for each model summarizing presence or absence of metabolites or reactions, observing that analyzed models were primarily grouped by database and method, inside of which phylogeny has greater influence, hence *C. griseus* and *A.aegypti* models were grouped together due to their closeness compared to *E. siliculosus*.

**Figure 5:**
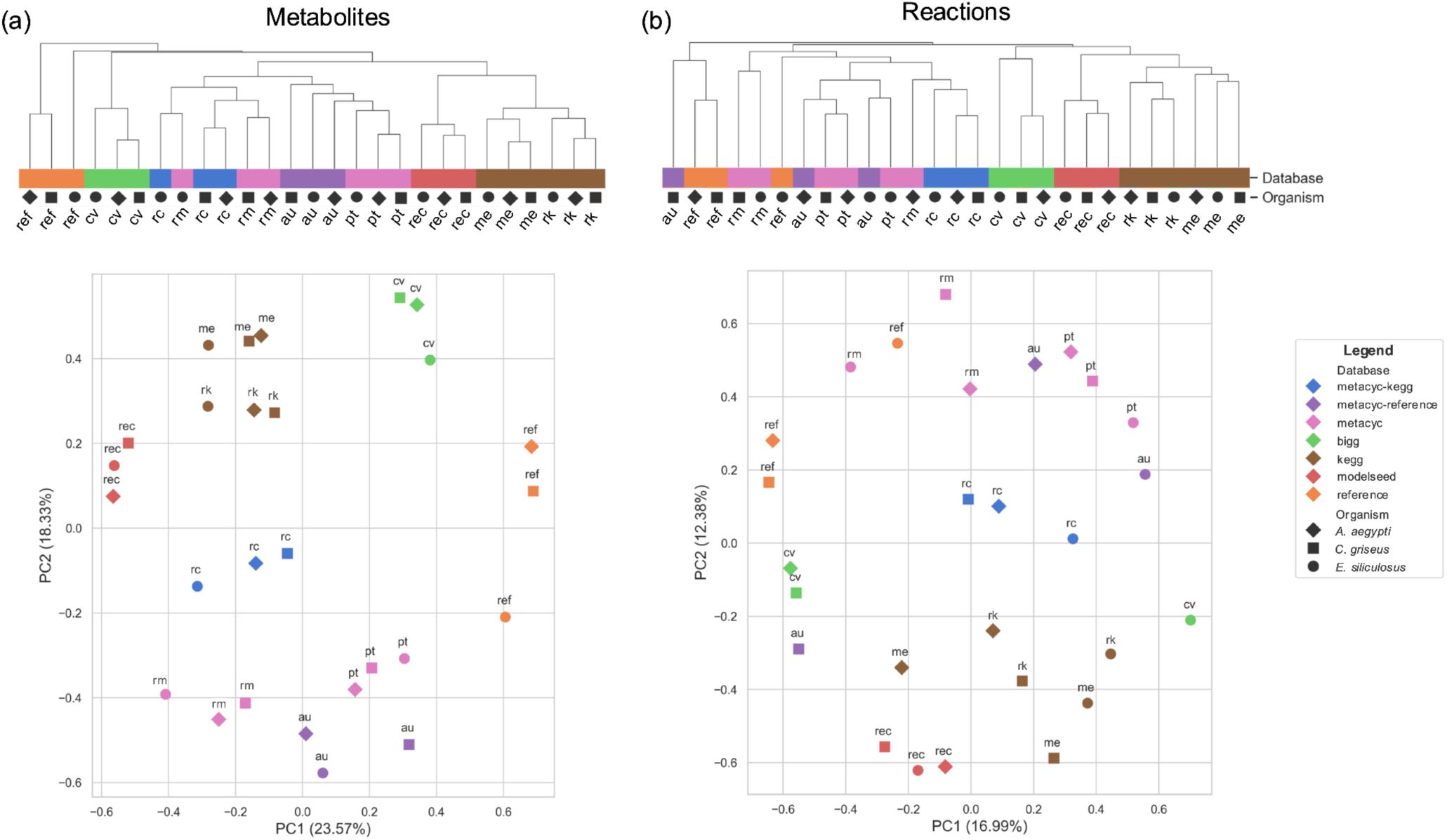
Influence of databases on metabolite and reaction inclusion in draft genome-scale models. Biclustering of reaction presence in draft models generated for this work, each method is presented with its associated database.

Models obtained using Reconstructor, Raven-kegg and Merlin, were clustered according to their associated databases (modelSEED and KEGG) in both metabolites and reaction analyses. Draft models generated using metacyc-based methods (AuReMe, pathway tools, raven-comb) are more similar with each other being clustered linked to their taxonomy, specifically with AuReMe, the only method where there is an intra-method disturbance in observed network structure given the influence of template databases, however they are similar to pathway tools drafts given the integration of an annotation step into their pipeline (**Supplementary File S8**).

*E. siliculosus* draft model was the only one found to be more similar to its respective Pathway tools draft model (Figure 5b), which could be associated with the choice of template model for its generation, the microalgae *Chlamydomonas reinhardtii*. In fact, the *C. griseus* model depicted here is based on a *Mus musculus* model (45), and the *A. aegypti* model is based on a *D. melanogaster* model (62), both of which are based on *H. sapiens* models.

A similar effect is observed for raven-metacyc and raven-comb models, where differences regarding the taxonomy of this macroalgae are more relevant that subtle differences observed in combined drafts obtained using this method.

Results are highly dependent on a good quality of mapping between databases, to achieve this goal metabolites were mapped into MetanetX, showing minimum mapping fractions of 96,41% for AuReMe-*E. siliculosus* (**Table S6**).

Due to the nature of this analysis, where included metabolite and reactions are compared between all the analyzed drafts; removal of compartment association is required. Hence, these results do not retrieve key differences associated with higher compartmentalization of these models. Methods like Merlin, that incorporate additional data of compartment associated reactions, are not depicted in this analysis; showing models from this method to be similar with drafts obtained using raven-kegg, which only includes one compartment on their models. Showing that although these analyses provide valuable insights on the information included in draft reconstructions, this constitutes a multidimensional problem that has to be approached from different perspectives.

## Discussion

Organism-specific reconstruction of GEMs is a complex process that can take up to years, which has motivated the development of automated tools that help accelerate this task. However, tool selection of which approach will be used is yet unclear and usually depends on the expertise of the team performing this task. An ideal model should be able to accurately represent the metabolic landscape of each organism, requiring the least amount of manual curation possible, while also being fit for typical GEM applications such as flux simulations and omics data integration. In this work we assessed seven methods in terms of their ability to represent organism- and compartment-specific metabolic features, ability to synthesize precursors for cell growth and annotation quality and how well they are annotated for their use in integration of omic data.

In this line, we selected metrics that represented an ideal draft model: one that includes the greatest number of reactions and unique metabolites (that is ignoring duplication of metabolites in several compartments), presents multiple compartments in lines with the analyzed organism, includes the greatest number of genes (enzymatic reactions), mass-balanced reactions (consistency score), how functional it is (biomass metabolites, flux reactions), and how well suited is for integration of omic datasets (annotation score). Some of these statistics are computed using Memote (42), a tool that proposes to set a standard for published GEMs.

Selected methods include tools originally developed for prokaryote organisms (carveMe, reconstructor) that have been adapted to their use in plants (plantSEED), which as expected displayed biomass representations based on gram positive or negative bacteria, as well as a reduced number of compartments. These methods were included based on their prevalence for their comparison with methods more compatible with eukaryotes, and although they retrieve in some cases closely to functional models their predictions should be used with caution given the phylogenetic distance between the organisms for which they were developed for in contrast with the organisms analyzed in this work.

One of the main challenges in the reconstruction process is the representation of organism-specific pathways, which are not always present in mostly automated databases. Methods based on manually curated databases like Metacyc, such as Pathway tools, Raven and AuReMe, pose an opportunity regarding representation of these pathways although, in general they do not retrieve functional models, or models characterized by higher percentage of unblocked reactions.

Accurate representation of subtle differences between compartment-specific enzymes is key for integration of transcriptomic, or proteomic datasets. In that regard, tools like Merlin, that perform an additional step for determining enzyme location, provide an effort in the right direction, providing a higher proportion of gene protein reaction associations on different compartments than other methods.

Other available methods, such as metaDraft (63) and gapseq (30), were considered for this comparison but were unable to retrieve any results for the analyzed organisms due mainly for technical issues associated with bigger genomes of these organisms, and lack of organism-specific data in KEGG (AutoKEGGRec, (23)).

As it was previously discussed for prokaryotes and unicellular eukaryotes (21), no method was universally better for generation of draft GEMs. Tool selection should depend on how well characterized the organism is, their intended application, and the expertise of the users, particularly their ability of fixing usual issues with boundary compartments and others that arise when installing and using tools developed by scientists in the systems biology field.

In particular, AuReMe is recommended for organisms that are not well documented in metabolic databases, given its ability to incorporate metabolic pathways added in manual curation steps for models from similar organisms. However the use of this approach has a steep learning curve, and does not result in functional models directly from its application.

If the intended use of the genome-scale model is to integrate high-throughput omic datasets (11,12) a method that specifically accounts for differences between compartments, such as Merlin, provides valuable insights that could significantly affect the obtained results. However, application of this tool is slow due to multiple steps required for its application.

Tools like carveMe and Reconstructor are characterized by retrieving close-to functional models in all the analyzed scenarios. Reconstructor seems to perform a less intensive gap-filling than carveMe, based on parsimony (22), which could be better suited for the study of organisms that are known to require interaction with their surrounding microbial community for growth and development (40). However, since both include composition of bacterial biomass into their models, an additional step for manual curation is required to remove the consequences of this assumption.

Given the staggering influence that databases have on the obtained drafts using different tools, we propose a preliminary exploration of the representation of metabolites of interest associated with the target organism to guide the selection of tools for draft generation. This difference in databases also affects analyses performed in this work, such as the one performed in analyzing presence of specialized metabolites, that were based on mapping metabolites into KEGG.

Systems biology is an interdisciplinary area of research that merges people with different backgrounds, the heterogeneity of this field results in applications of genome-scale models from exploration of metabolic capabilities (34), biotechnological applications (35,64,65) and even analysis with ecological implications (66). Hence, the need for tools with different approaches designed with different applications in mind. In this work we provided thorough analyses that attempt to provide light on the selection of tools for generation of genome-scale models in multicellular eukaryotic organisms

## Materials and methods

### Draft generation

Three organisms from a wide range of phylogenetic groups and different degrees of knowledge were selected to make this evaluation: (i) *Cricetulus griseus* and its derived cell line CHO (Chinese Hamster Ovary), for which there is a highly curated genome and GEM (35) and has been widely used in different applications (19,67), (ii) *Ectocarpus siliculosus*, a brown kelp for which there is a genome-scale model (34), and (iii) *Aedes aegypti*, a mosquito for which there is a recent metabolic reconstruction (36).

*Cricetulus griseus* genome (CriGri_1.0) (68) and its annotation were retrieved from NCBI (GCA_000223135.1). *A. aegypti* genome (AaegL5.0) (Matthews et al., 2018) and its annotation were retrieved from vectorBase. *E. siliculosus* genome (ASM31002v1) (70) and its annotation were retrieved from NCBI (GCA_000310025.1).

Seven frameworks for generation of draft genome-scale models were retrieved based on literature and their capabilities to handle eukaryotic genomes. The process to generate draft models using these approaches are described below.

### AuReMe

Annotation and orthology-based drafts were generated using AuReMe (version 2.4) as follows: organism-specific Pathway tools annotation folders (.dat folder) were provided for the annotation-based step, and different reference organisms with their respective models were used for the analyzed organisms in this work. *Drosophila melanogaster* (Release 6 plus ISO1 MT, GCF_000001215.4) was used as reference organism for *A. aegypti* (71), *Mus musculus* (GRCm39, GCF_000001635.27) (72) for CHO (*C. griseus*) (45) and *Chlamydomonas reinhardtii* (Chlamydomonas_reinhardtii_v5.5, GCF_000002595.2) (73) for *E. siliculosus* (44). Additional reconstructions were obtained using *Homo sapiens* (GRCh38.p14, GCF_000001405.40) (46) for *A. aegypti* and CHO, and *Saccharina japonica* (31,43) for *E. siliculosus* to test the effect of reference organisms in the obtained quality of reconstructions.

### CarveMe

CarveMe (27) was used with amino acid fasta files of the target organisms as input using their default settings (bacterial template and the arguments --more-sensitive --top 10 for DIAMOND (74)).

### Merlin

Merlin 4.014 (25) was used, homology search was performed using diamond (74) with the default e-value (1e-30) and KEGG metabolic data was loaded into their workspace. Enzyme annotation and integration were performed through their automatic workflow complemented by Transyt (75) to create transport reactions. Subsequently, the compartment report from Deeploc2 (48) was loaded and integrated into the model. Finally, the biomass equation was created using yeast as a template.

### ModelSeed

An implementation of modelSEED for eukaryote organisms (plantSEED) (29) was used with amino acid fasta files of the target organisms studied. A ‘complete’ medium was selected to perform draft model generation, which considers presence of a metabolite in the media only if there are transporters for this metabolite in the model.

### Pathway tools

Genome annotation was complemented with additional data retrieved from genome annotation using eggnogmapper (76) with their web based version using standard parameters (minimum hit e-value: 0.001, Minimum hit bit-score: 60, percentage identity: 40, minimum % of query coverage: 20, minimum % of subject coverage: 20), using Eukaryota as their taxonomic scope. Obtained results were then combined to generate gbk files using emapper2gbk (77) to retrieve gbk files compatible with Pathway tools, which were subsequently used to generate the draft models with default parameters.

### Raven 2

Implementation of Raven 2 (Matlab 2015a) was used to generate a combined draft GEM from KEGG and Metacyc as specified on their documentation (24). In summary, a metacyc-based draft (‘raven-metacyc’ in this work) was generated from amino acid fasta files, keeping transport reactions with other parameters set as default (exclusion of unbalanced and undetermined reactions, minimum bit score: 100, minimum positive values: 45%, diamond as alignment tool to perform homology search). A KEGG-draft was generated based on sequence homology (‘raven-kegg’) of fasta amino acids file to KEGG Ortholog sequence clusters, incorporating the eukaryote cluster “euk90_kegg105”, while excluding incomplete reactions and the ones with undefined stoichiometry, the other parameters are set as default (cutOff: 1e-50, minScoreRatioG: 0.8, minScoreRatioKO: 0.3, seqIdentity=0.9)

Finally both draft models were combined into an integrated GEM (‘raven-comb’) using the function combineMetaCycKEGGModels, which converts metabolite and reaction identifiers in the KEGG model into corresponding MetaCyc IDs, and then detects duplications and keeps only unique reactions and metabolites that are mostly in the MetaCyc namespace.

### Reconstructor

Reconstructor (22) was used with amino acid fasta files of the target organisms as input, using default parameters (minimum objective fraction required during gap filling: 0.01, maximum objective fraction allowed during gap filling: 0.5) with a Gram-negative classification.

### Analysis of draft models

Different metrics were selected to assess features of the obtained draft models, memote (42) was used to compute the number of genes, reactions, metabolites and compartments, as well as gene associated reactions (enzymatic reactions) and reactions able to carry flux (flux reactions) of the obtained models in complete medium. Execution times were logged manually for their analysis.

### Metabolite and reaction mapping

Metabolites and reactions from different namespaces were translated to MetaNetX namespace (Moretti et al., 2021) for comparing model contents. Two different mapping strategies were used: (i) direct mapping via the MetaNetX database, after removing compartment-related suffixes for deduplication of biochemically equivalent reactions. This procedure was complemented with information from BiGG (79), modelSEED (28) and the KEGG (80) REST API regarding metabolite names and chemical composition. Stoichiometry was used as additional information for reaction mapping as described previously (21). (ii) Model mapping using Python package mergem v.1.1.0 (Hari et al. 2024), with MetaNetX as the target database. Compartment-related suffixes were removed after mergem mapping; manual curation of the obtained mappings was performed to resolve conflicting mappings between these alternative methods.

### Biomass production

Biomass composition was retrieved from published models for *A. aegypti* (36), *C. griseus* (35) and *E. siliculosus* (34) and translated into other databases as needed (Metacyc (81), bigg (79), modelSEED, KEGG (80)). Draft models were scored according to the extent of inclusion and producibility of these expected metabolites. A score 0 was assigned to absent metabolites, a score 1 was assigned to metabolites that are present but not producible and a score of 2 was assigned to present and producible metabolites Producibility was assessed via FBA simulations using COBRApy (82) with a complete medium obtained by opening all exchanges of the model and setting the production of each metabolite as the objective function. A final score for each model was computed as the sum of all metabolite scores divided by the maximum score possible for each biomass function. Metabolites were grouped into categories for its visualization using MetaboAnalyst (83) followed by manual curation.

### Mapping specialized metabolites

Organism specific metabolome data was retrieved from literature for *Aedes aegypti* (50), CHO cells (35,51–59) and *E. siliculosus* (60). Raw data was mapped to KEGG identifiers using metaboanalyst 6.0 (84), presence of mapped metabolites was assessed for all generated drafts in the KEGG namespace.

### Compartment analysis

Duplicated reactions were selected based on the participation of duplicated metabolites and their stoichiometry. For each duplicated reaction, gene associations were studied to determine if they were identical to the cytoplasmic form of that reaction, and classified in terms of presence of GPR and their uniqueness. Plots were made using seaborn (85,86).

### Model comparison

Obtained drafts were compared in regards to their genes, reactions and metabolites. For reaction and metabolite analyses, a binary representation was constructed for each model as follows: for each metabolite or reaction a number 1 was assigned if it is present in the model. All models were mapped to the MetaNetX database (78) as previously described

To study the effects of the reference models in AuReMe, three models were used for comparison: one based solely on annotation, and two combining this reconstruction with orthology—the one used in previous analyses (see AuReMe section in methods), and another employing additional reference models *H. sapiens* was used as the reference for *A. aegypti* and *C. griseus* (46), and *S. japonica* for *E. siliculosus* (43). Results were compared at metabolite, reaction and gene levels, by extracting this information using COBRApy (82), and plotted using matplotlib (85). Due to the presence of metabolites with identifiers from different databases, the script described in the “Metabolite and Reaction Mapping” section was used to ensure proper correspondence.

An additional analysis, where genes are compared to their reference genomes, was performed by mapping gene identifiers from models to their genome using SynGO and bioDB mapping tools (87,88). For *C. griseus* DIAMOND was used to retrieve equivalences between RefSeq and GenBank where direct mapping was unavailable, selecting the match with the highest bit score.

## Acknowledgements

This research was funded by ANID, Núcleo Milenio MASH NCN2021033, NEMBICA STIC190013, and Fondecyt Iniciación 11090268.

NEJ, IS, JCS, CC and ZGH received support from ANID CeBiB PFBasal-0001, ZGH and JCS from ANID FONDEF IT15I10048, SM and CC from CMM FB210005.

## Author contributions Statement

Authors CC, JCS, NEJ and ZGH worked on the conceptualization of this study. Data acquisition was carried out by IS and ME, analysis was performed by IS, ME, NEJ and SM. Interpretation of obtained results was achieved by CC, IS, JCS, ME, NEJ, SM, and ZGH. IS, ME and NEJ wrote the first draft of this work, which was revised by all authors prior to submission.

## Availability of data and Materials

Data (obtained models as well as their analysis) is provided within the manuscript, supplementary information files or the github repository https://github.com/natJimenez/eukaryo_methods.

## Supplementary material

- **S1 Table**: Metrics of generated draft genome-scale models
- **S2 Table**: Analysis of gene distribution in plantSEED generated models
- **S3 Figure**: Description of biomass composition for reference models
- **S4 Table** Metabolite list for metabolome mapping
- **S5 Table**: Standard deviation of metabolite mapping of specialized metabolite datasets
- **S6 Table**: Mapping percentages for metabolites and reactions
- **S7 Figure**: Mapping reactions and metabolites
- **S8 File**: Influence of template model in AuReMe generated drafts

## S8 - Effect of template model in AuReMe

Further analysis with alternative template models using this tool showed that predictions made from orthology exhibit a trade off between phylogenetic closeness and the information contained in models used for their reconstruction (**Figure S8.1**). Phylogenetic distance within modeled organisms is a key differentiating factor at a metabolic level for both metabolite and reaction content included in the analyzed drafts, where models for *E. siliculosus* are grouped together at both analyzed levels with the exception of the draft retrieved from orthology between *E. siliculosus* and *C. reinhardtii*. This model retrieves reactions linked to the orthology step for our original template selected for this organism, a microalgae which could explain their behaviour at both levels. On the other hand, template models from phylogenetically close organisms *(Mus musculus* and *Homo sapiens* for *C. griseus* draft generation) retrieve almost identical models at a metabolite level and similar models at a reaction level.

**Figure S8.1:**
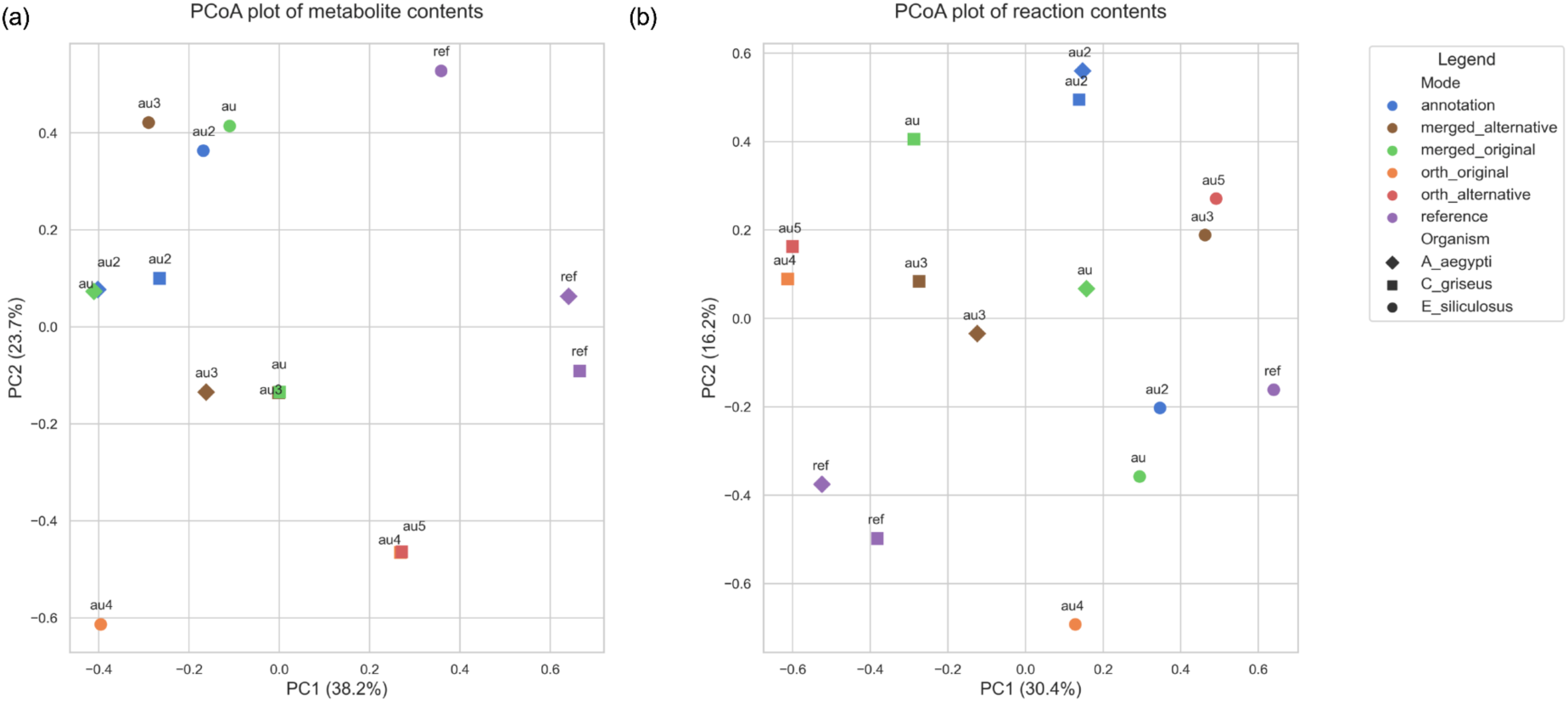
Effect of different inputs and stages of generation in AuReMe draft generation models on metabolite (a) and reaction (b) contents. From annotation steps performed by Pathway tools (annotation), orthology steps performed using original templates (orth_original) as well as alternative templates (orth_alternative) as specified in methods; to finally, merged models using both approaches (merged_original, merged_alternative). Reference models are included to depict their similarity to obtained draft models.

Similarities between different stages of draft generation using the AuReMe workflow (annotation, orthology, merged) are observed. For *Ectocarpus*, merged and annotation models from original template are more similar with each other than with orthology template, alternative template (*S. japonica*) retrieves a model which is closer with these models, hinting a possible database-specific effect. For *Cricetulus* and *Aedes* the effect of the alternative template was strikingly clear, with the *Aedes* alternative draft model (template: *H. sapiens*) was more similar to the *C. griseus* draft based on the same template model.

